# Tissue-specific volatile-mediated defense regulation in maize leaves and roots

**DOI:** 10.1101/2020.02.21.959437

**Authors:** Cong van Doan, Tobias Züst, Corina Maurer, Xi Zhang, Ricardo A.R. Machado, Pierre Mateo, Meng Ye, Bernardus C.J. Schimmel, Gaétan Glauser, Christelle A.M. Robert

**Affiliations:** Institute of Plant Sciences, University of Bern, Bern, Switzerland; Oeschger Centre for Climate Change Research (OCCR), University of Bern, Bern, Switzerland; Key Laboratory of Plant Stress Biology, State Key Laboratory of Cotton Biology, School of Life Sciences, Henan University, Kaifeng 475004, China; Neuchâtel Platform of Analytical Chemistry, Université de Neuchâtel, Neuchâtel, Switzerland

**Keywords:** belowground plant-herbivore interactions, maize, plant-plant interactions, priming, volatiles

## Abstract

- Plant leaves that are exposed to herbivore induced plant volatiles (HIPVs) respond by increasing their defenses. Whether this phenomenon also occurs in the roots is unknown.
- Using maize (*Zea mays*), whose leaves respond strongly to leaf HIPVs, we measured the impact of root HIPVs, emanating from plants infested by the banded cucumber beetle (*Diabrotica balteata*), on constitutive and herbivore-induced levels of root soluble sugars, starch, total soluble proteins, free amino acids, volatile and non-volatile secondary metabolites, defense gene expression, growth and root herbivore resistance of neighboring plants.
- HIPV exposure did not alter constitutive or induced levels of any of the measured root traits. Furthermore, HIPV exposure did not reduce the performance and survival of banded cucumber beetle larvae on maize or teosinte. Cross-exposure experiments revealed that maize roots, in contrast to maize leaves, neither emit nor respond strongly to defense-regulating HIPVs.
- Together, these results demonstrate that volatile-mediated defense regulation is restricted to the leaves of maize and teosinte, a finding which is in line with the lower diffusibility of volatiles in the soil and the availability of other, potentially more efficient information conduits below ground.

## INTRODUCTION

Upon herbivory, plants emit volatile organic compounds that can repel herbivores and attract their natural enemies (Baldwin, 2010; Turlings & Erb, 2018). These herbivore-induced plant volatiles (HIPVs) can also be perceived by unattacked plant tissues and neighboring plants, resulting in the direct activation and/or priming of defense and resistance (Farmer, 2001; Baldwin *et al*., 2006; Frost *et al*., 2008; Heil & Ton, 2008; Heil, 2014; Erb, 2018; Turlings & Erb, 2018; Bouwmeester *et al*., 2019). Numerous HIPVs have been found to regulate defenses, including green leaf volatiles such as (Z)‐3‐ hexenal, (Z)‐3‐hexen‐1‐ol, and (Z)‐3‐hexenyl acetate (HAC), aromatic compounds such as indole, and terpenoids such as ocimene (Farmer, 2001; Engelberth *et al*., 2004; Erb *et al*., 2015; Riedlmeier *et al*., 2017; Ameye *et al*., 2018). HIPVs can regulate redox signalling genes (González-Bosch, 2018), early defense signalling genes and proteins such as MAP kinases (Ton *et al*., 2007; Erb *et al*., 2015; Hu *et al*., 2019; Ye *et al*., 2019), the biosynthesis of stress hormones such as jasmonates (Ton *et al*., 2007; Heil & Ton, 2008; Hirao *et al*., 2012) and the expression of direct and indirect defenses (Zeringue, 1987; Zeringue, 1992; Bate & Rothstein, 1998; Arimura *et al*., 2000; Arimura *et al*., 2001; Engelberth *et al*., 2004; Farag *et al*., 2005; Kessler *et al*., 2006; Kost & Heil, 2006; Ton *et al*., 2006; Karban, 2011; Kim *et al*., 2011; Erb *et al*., 2015; Martinez-Medina *et al*., 2016; Freundlich & Frost, 2018; Tugizimana *et al*., 2018).

Although defense regulation by HIPVs has been documented extensively in plant leaves, much less is known about this phenomenon in the roots (Delory *et al*., 2016). To the best of our knowledge, no study so far investigated the impact of root HIPVs on defense and resistance of neighboring plants. Roots emit specific volatile blends when attacked by herbivores (Rasmann *et al*., 2005; Ali *et al*., 2010; Delory *et al*., 2016). These volatiles can diffuse through the soil and alter the behaviour of herbivores and natural enemies (Hiltpold & Turlings, 2008; Xavier *et al*., 2017; Gfeller *et al*., 2019). Recent work also found that constitutively released root volatiles can affect growth and defense expression in neighboring plants (Huang *et al*., 2018; Gfeller *et al*., 2019). Thus, it is conceivable that roots may also respond to root HIPVs in anticipation of root herbivore attack.

To test this hypothesis, we investigated HIPV-mediated root interactions in maize, one of the three most important crops worldwide (Shiferaw *et al*., 2011). Maize plants are regularly attacked by root herbivores such as rootworms, which can cause substantial damage and yield losses (Tinsley *et al*., 2016). Maize leaves are highly responsive to leaf HIPVs such as indole and (*Z*)-3-hexenyl acetate (Engelberth *et al*., 2004; Erb *et al*., 2015; Hu *et al*., 2019). Upon herbivore attack, maize roots emit distinct blends of HIPVs that contain terpenes such (*E*)-β-caryophyllene, humulene and copaene (Rasmann *et al*., 2005; Robert *et al*., 2012b; Robert *et al*., 2012a), but no detectable amounts of indole or GLVs. (*E*)-β-caryophyllene can diffuse up to 20 cm.h^−1^ in the soil matrix (Xavier *et al*., 2017). To test if maize roots can use root HIPVs to prepare their defense system for incoming herbivore attack, we first assessed the impact of root HIPVs on maize primary metabolism and defense markers in the absence of herbivory. Second, we assessed the impact of root HIPVs on root-herbivory induced changes in primary metabolism and defense markers. Third, we tested the effect HIPVs on plant growth and resistance. Fourth, we conduced cross-exposure experiments to assess the impact of leaf HIPVs on root resistance and *vice versa*. These experiments found no evidence for HIPV-mediated induction of root defenses, and suggest that roots do not respond to HIPVs by increasing their resistance to herbivores.

## MATERIALS AND METHODS

### Plants and insects

Maize seeds (*Zea mays* L., var. “Delprim”) were provided by Delley Semences et Plantes (DSP, Delley, CHE). Maize seeds were sown in plastic pots (diameter, 4cm; height, 11.2 cm; Patz GmbH Medizintechnik, Dorsten-Wulfen; DE) as described in (Erb *et al*., 2011). The seedlings were fertilized twice a week after germination with MioPlant Vegetal and Herbal Fertilizer (Migros, CHE). Twelve-day old plants with three fully developed leaves were used for the experiments. Eggs of the banded cucumber beetle *Diabrotica balteata* (Coleoptera: Chrysomelidae) were kindly provided by Oliver Kindler (Syngenta, Stein, CHE). Hatching larvae were reared on freshly germinated maize seedlings (var. Akku, DSP, CHE). Second-instar larvae were used in the experiments. The larval instars were determined according to the head capsule size as previously described (George & Hintz, 1966). Plant infestations were performed by placing six larvae in two 4-5 cm deep holes in the sand. Eggs of the Egyptian cotton leafworm *Spodoptera littoralis* were provided by the University of Neuchâtel and reared on artificial diet until use.

### Characterization of root HIPV production by emitter plants

To determine the HIPV profile emitted by root-infested plants over time, maize plants were placed into L-shaped glass pots (diameter: 5 cm; depth: 11 cm; Verre & Quartz Technique SA, Neuchâtel, CHE). Moist white sand (Migros, CHE) was added to fill the pots. The L-pots were wrapped in aluminium foil to keep the root system in the dark and prevent degradation of volatile compounds. Two days later, half the plants were infested with six second-instar *D. balteata* larvae. Control and infested maize roots were collected after one, two, three, four or eight days (n=5-7 per treatment and per day). The roots were ground in liquid nitrogen using a mortar and a pestle. An aliquot of 100 mg was used to measure root volatile production by solid phase micro extraction gas chromatography coupled to mass spectrometry (SPME-GC-MS, Agilent 7820A GC coupled to an Agilent 5977E MS, Agilent Technologies, Santa Clara, CA, USA). Briefly, a 100 µm polydimethylsiloxane SPME fibre (Supelco, Bellefonte, PA, USA) was inserted through the septum of the root containing glass vial (20 mL Precision Thread Headspace-Vial and UltraClean 18 mm Screw caps, Gerstel GmbH & Co., Mülheim an der Ruhr, DE) and exposed to the vial headspace for 40 min at 50°C. The fibre was inserted into the GC injection port (220°C) and desorbed. Chromatography was performed using an apolar column (DB1‐MS, 30 m, 0.25 mm internal diameter, 0.25 µm film thickness; J & W Scientific, Folsom, CA, USA). Helium was used as carrier gas at a constant pressure of 50.6 kPa. The column temperature was maintained at 60 °C for 1 min and then increased to 250 °C at 5 °C min^−1^ followed by a final stage of 4 min at 250 °C. Volatile identification was obtained by comparing their mass spectra with those of the NIST05 Mass Spectra Library.

### Root herbivore migration timing

To determine the most realistic experimental timing for the response phase of neighboring plants, we evaluated the time window during which *D. balteata* root herbivores are most likely to migrate from an infested to a neighboring plant. Maize plants were potted into 100 mL pots with two 5 mm diameter openings at the bottom. Each pot was placed in a plastic cup (12 × 25 × 10 cm WxLxH, OBI Group Holding SE & Co.KGaA, Schaffhausen, CHE) filled with a 3 cm high layer of tap water. All plants (n=6) were infested with six second-instar *D. balteata* larvae. The larvae moving away from the plant through the openings or from the top of the pot were therefore trapped in water and collected daily.

### Exposure to belowground HIPVs

To test whether plant exposure to belowground HIPVs induces a response in neighboring plants, belowground two-arm olfactometers were used as previously described (Robert *et al*., 2012a). Briefly, maize plants were placed into L-shaped glass pots (diameter: 5 cm; depth: 11 cm). Moist white sand (Migros, CHE) was added to fill the pots. The L-pots were wrapped in aluminium foil to keep the root system in the dark and prevent degradation of volatile compounds. Two days later, pots containing plants of similar sizes were connected in pairs using two Teflon connectors and one glass connector (length, 8 cm; diameter, 2.2 cm, VQT, Neuchâtel, CHE). The Teflon connectors contained a fine metal screen (2300 mesh; Small Parts Inc., Miami Lakes, FL, USA) to restrain the larvae from moving to the second plant. The glass connectors remained empty to only allow volatile compounds to diffuse through the system. Each pair included one emitter plant and one receiver plant. Emitter plants were either infested with six second-instar *D. balteata* larvae or remained uninfested. Receiver plants were exposed to emitter plants for four days prior to any treatment. After this four days exposure period, receiver plants were either infested with six root herbivore larvae or left uninfested depending on the experiments. All pairs remained connected until collection of the samples.

### Root responses to root HIPVs

To evaluate how exposure to HIPVs affects the metabolism of maize plants in absence and presence of herbivores, two independent experiments were conducted. In the first experiment, primary metabolism and defenses of receiver plants were characterized after four days exposure to HIPVs in absence of herbivory (n=9 per treatment). In the second experiment, receiver plants were infested with six second-instar *D. balteata* larvae, and primary metabolism and defenses were measured 1, 3, 6, 9 and 12 hr after the onset of herbivory (n=3-7). In all experiments, maize roots were collected, gently washed with tap water, flash frozen in liquid nitrogen and ground to a fine powder for further analyses. Plant primary metabolism was assessed by measuring sucrose, glucose, fructose and starch using enzymatic assays (Velterop & Vos, 2001; Smith & Zeeman, 2006; Machado *et al*., 2013), soluble proteins using colorimetric assays (Bradford, 1976; Jongsma *et al*., 1994), free amino acids using HPLC-MS (Li *et al*., 2018), and the expression of the carbohydrate transporters *Zm-stp1, Zm-zifl2* by q-RT-PCR (Robert *et al*., 2012b) (Supporting Information Table S1). Plant secondary metabolism was characterized by performing untargeted metabolomic analyses by UHPLC-qTOF-MS (Hu *et al*., 2018), measuring concentrations of benzoxazinoids by UHPLC-qTOF-MS (Hu *et al*., 2018), and volatile emissions by GC-MS as described above. Plant defense expression was characterized by measuring stress hormones by UHPLC-MS/MS (Glauser *et al*., 2014) and defense marker genes, including genes involved in volatile production (*Zm-tps23, Zm-igl*),; hormonal signalling (*Zm-saur2, Zm-nced, Zm-orp7, Zm-lox5 Zm-acs6*) and direct defenses (*Zm-cysII, Zm-cyst, Zm-serpin, Zm-mpi, Zm-bx1, Zm-pal, Zm-pr1*) by q-RT-PCR (Robert *et al*. 2012b). For a more detailed description of these genes, refer to (Robert *et al*. 2012b) and Supplementary Information Table S1.

### Plant and herbivore performance following root exposure to root HIPVs

To determine whether exposure to root HIPVs impacts the performance of root herbivores, belowground two-arm olfactometers were used as described above. After four days exposure to control or infested emitter plants, all receiver plants were infested with six pre-weighed root herbivore larvae (n=18 per treatment). Four days later, all larvae feeding on receiver plants were recovered and weighed. Maize roots from the plants were collected for damage evaluation (Oleson *et al*., 2005) and weighed.

### Cross-exposure experiment

To assess whether priming is tissue-specific, cross exposure experiments were conducted by exposing roots or leaves to volatiles emitted by either control or infested roots or leaves of emitter plants (n=4-5 per treatment). All plants were potted in L-pots as described above. Emitter plants were either infested with six second-instar *D. balteata* (root herbivory), three fourth-instar *S. littoralis* larvae (leaf herbivory) or left uninfested. All plants were covered with plastic bags (Bratbeutel Tangan N°34, Genossenschaft Migros Aare, Urtenen-Schönbühl, CHE). Emitter and receiver plants were paired using the glass connectors described above. The glass connectors were used to connect roots to roots, roots to leaves, leaves to roots or leaves to leaves. To connect a leaf compartment, a 3 cm opening was made in the plastic bag to insert the connector. The bag was then sealed around the glass connector with a rubber band and tape. The headspace of emitter plants was connected to a multiple air-delivery system via PTFE tubing. Purified air was pushed in the system at a flow rate of 0.3 L.min^−1^. After 17 hr exposure to emitter plants (from 5 pm to 10 am the next day), all systems were disconnected and bags removed. Three pre-weighed *S. littoralis* or six pre-weighed second-instar *D. balteata* larvae were added to receiver plants and new plastic bags were added to all plants. After 2 days, all larvae were collected and weighed.

### Statistical Analyses

Statistical analyses were conducted using R (version 3.5.3, https://www.r-project.org) and Sigma Plot (version 13, Systat Software, San Jose, CA). All data were first tested for normality and heteroscedasticity of error variance using Shapiro-Wilk and Brown-Forsythe tests. Data fitting normality and variance equality assumptions were analyzed using Analysis of Variance (ANOVA). Data that did not fit normality and equality of variance were analyzed using Mann-Whitney Rank Sum tests (U tests) and ANOVAs on ranks. Metabolomic and volatile data were analyzed using principal component analyses (PCA) followed by PPLS-DA and permutation tests.

## RESULTS

### Root herbivory induces a distinct bouquet of root volatiles

To characterize belowground HIPVs, we measured root volatile production from the plants over 8 days infestation. Root-herbivore infested plants produced distinct bouquets of volatile compounds over the entire exposure period, including high amounts of *(E)*-β-caryophyllene, caryophyllene oxide and copaene (Fig. 1).

**Figure 1.**
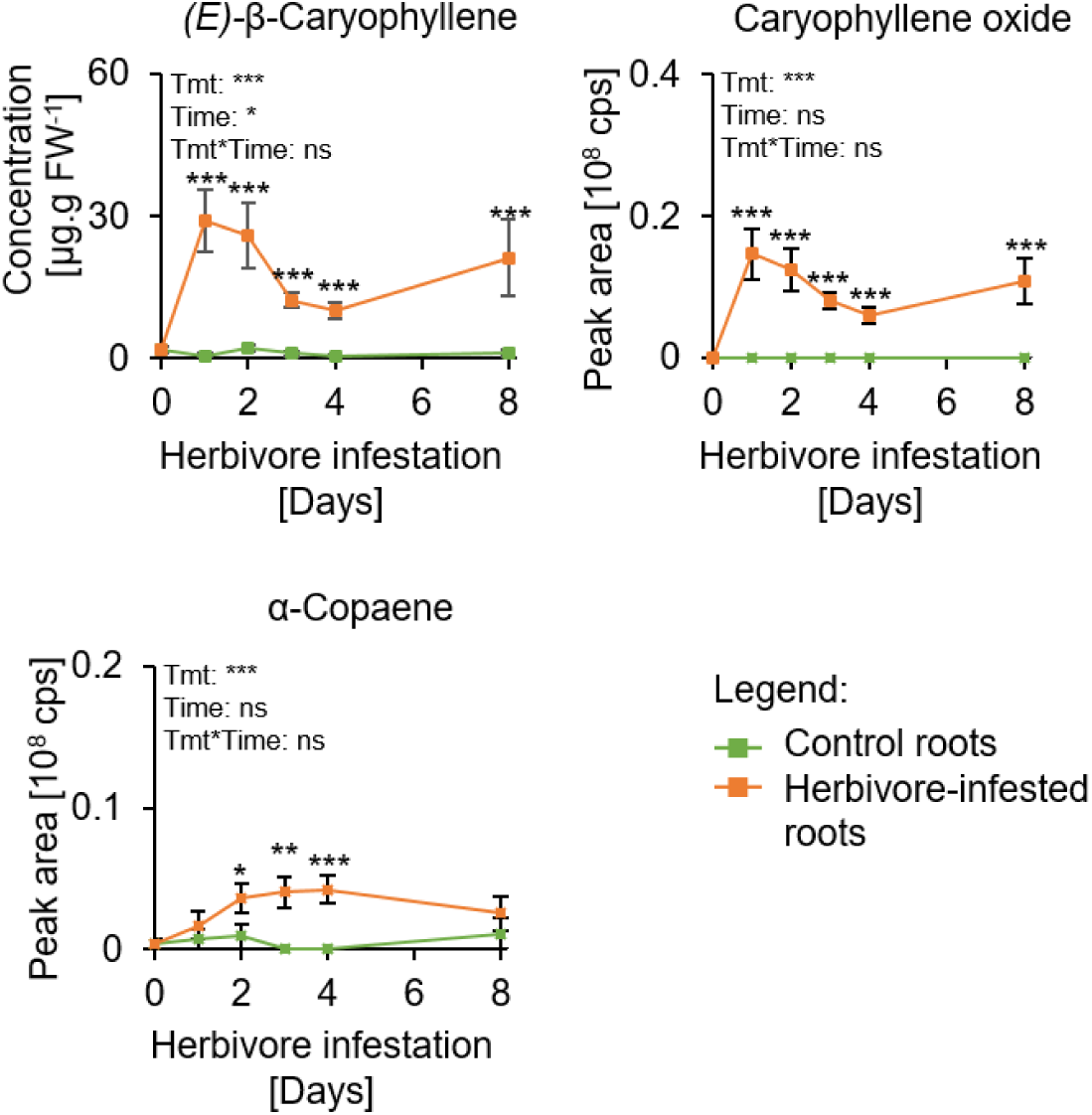
Root herbivory induces terpene volatiles from maize root. *(E)-*β-caryophyllene, caryophyllene oxide, and α-copaene emissions by control (green) and infested maize roots (orange) after 0-8 days (Mean ± se, Two way ANOVA, n=5-7). *(E)-*β-Caryophyllene was identified and quantified using a standard curve of the pure compound. Caryophyllene oxide and α-copaene were identified by using the NIST library (Match >85%). Tmt: Treatment. cps: Counts per second. Stars indicate significant differences (*: p≤0.05).

### Root herbivores migrate away from infested plants 1-4 days after the start of infestation

To assess the probability of a neighboring plant to be attacked, we measured larval migration from the plants over time. Root herbivore larvae migrated away from the first day on: After one day, 23.3% of the larvae were recovered outside the pots, and after four days, more than 60% had migrated away from the plant (Supplementary Information Fig. S1). Thus, response plants were exposed to root HIPVs for four days in subsequent experiments.

### Root HIPVs do not directly induce defenses in neighboring root systems

To evaluate whether belowground exposure to root HIPVs induces physiological changes in neighboring plants, we characterized the primary metabolism and defenses of maize roots exposed to control or root-herbivore infested volatiles over four days. The expression of marker genes involved in plant primary and secondary metabolism was not significantly altered by HIPV exposure (Fig. 2a). Phytohormone production was similar between control and HIPV-exposed roots, except for jasmonic acid (JA) and its isoleucine conjugate (JA-Ile), for which levels were slightly lower in HIPV-exposed roots than control roots (Fig. 2b). Individual and total soluble sugars, starch, protein, and amino acid concentrations were not affected by exposure to root HIPVs (Figs. 2c-e). Also, no significant effects on benzoxazinoids, the most abundant root secondary metabolites, were observed (Fig. 2f). Untargeted metabolomics (511 and 1763 features were detected in negative and positive modes, respectively) did not reveal differential clustering of chemicals (Figs. 2h-i). Finally, root volatile production remained unchanged between control and HIPV-exposed plants (Figs. 2g and j).

**Figure 2.**
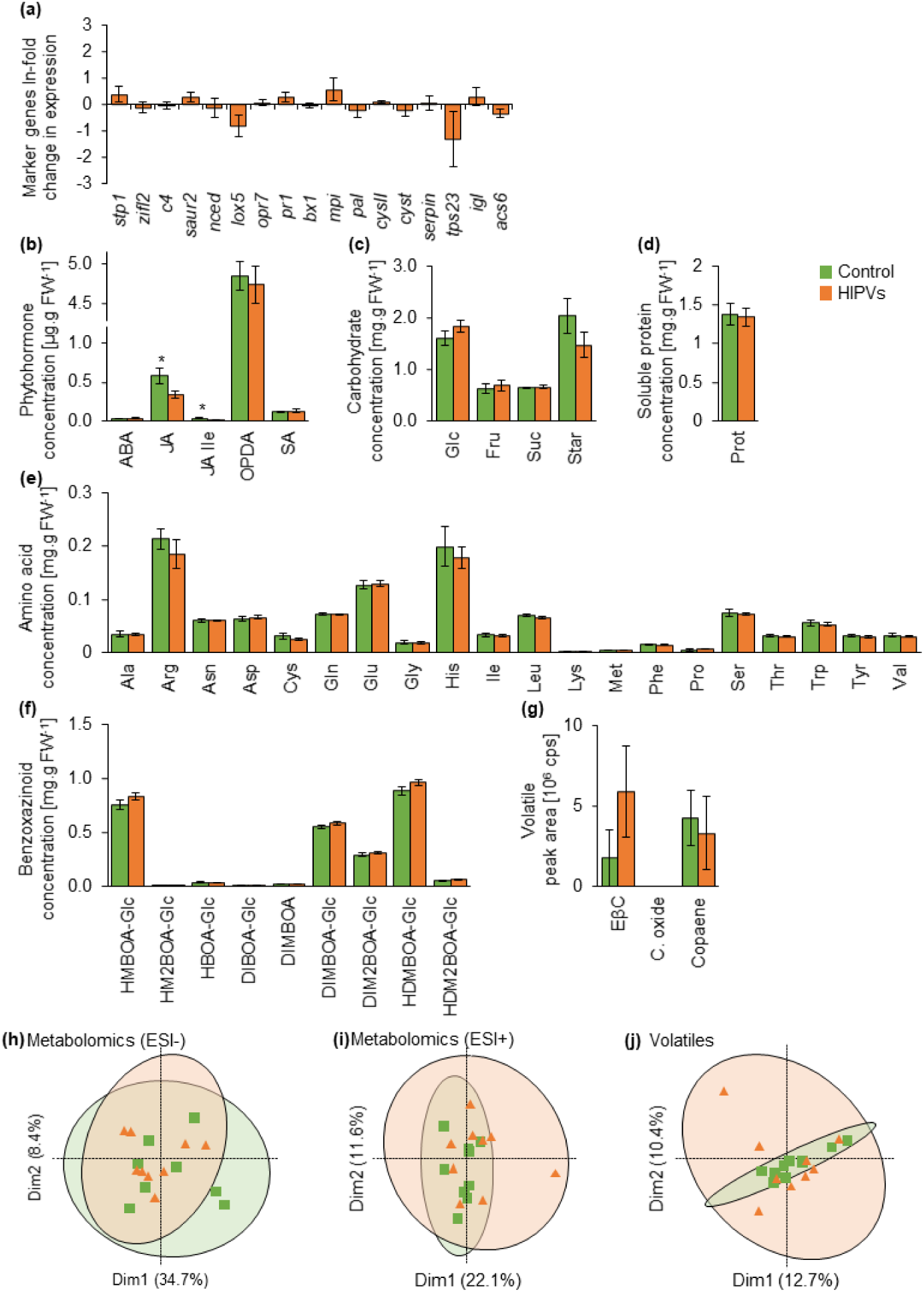
Belowground herbivore-induced plant volatiles (HIPVs) do not affect plant metabolism in absence of herbivory. (**a**) Ln fold changes in gene expression (Mean ± se, Student’s t-tests and Mann-Whitney U tests, n =9) in maize roots exposed for four days to plants infested with six *Diabrotica balteata* larvae (HIPVs) relative to maize roots exposed to control plants. (**b**) Phytohormone production (Mean ± se, Mann-Whitney U tests, n = 9) in maize roots exposed for four days to control plants (control, green) or to plants infested with six *D. balteata* larvae (HIPVs, orange). (**c-f**) Concentrations (Mean ± se, Student’s t-tests and Mann-Whitney U tests, n = 9) of (**c**) glucose, fructose, sucrose, and starch, (**d**) proteins, (**e**) amino acids, and (**f**) benzoxazinoids in roots of maize plants exposed for four days to control plants (control, green) or to plants infested with six *D. balteata* larvae (HIPVs, orange). (**h-i**) Principal Component Analysis of all features detected (PPLS DA, n = 9) in roots of maize plants exposed for four days to control plants (control, green) or to plants infested with six *D. balteata* larvae (HIPVs, orange) using untargeted metabolomic analysis in (**h**) negative (511 features) and (**i**) positive modes (1763 features). (**j**) Principal Component Analysis of volatile emissions (PPLS DA, n = 9) and (**g**) terpene volatiles emissions by roots of maize plants exposed for four days to control plants (control, green) or to plants infested with six *D. balteata* larvae (HIPVs, orange). EβC: *(E)-*β-caryophyllene. C. oxide: Caryophyllene oxide. Stars indicate significant differences (*: p≤0.05).

### Root HIPVs do not change root defense induction in neighboring root systems

To investigate whether belowground HIPV-exposure alters responses to herbivory in the roots of neighboring plants, we characterized root responses to infestation by *D. balteata*. Marker genes involved in plant response to root herbivory (Robert *et al*., 2012b) responded similarly in control and HIPV-exposed maize plants, with the exception of *acs6* (Fig. 3a). The production of abscisic acid (ABA), oxo-phytodienoic acid (OPDA) and JA and JA-Ile increased upon root herbivory but was not influenced by HIPV exposure (Fig. 3b). Carbohydrate concentrations were similar in control than in HIPV-exposed plants although HIPV-exposed plants overall had lower fructose concentrations than control plants (Fig. 3c). Soluble proteins, and amino acids responded to herbivory independently of HIPV exposure (Figs. 3d-e). Untargeted metabolomics (443 and 1906 features detected in negative and positive modes, respectively) and benzoxazinoid profiling did not reveal differential clustering or differences in concentrations (Figs. 3f, h-i). Volatiles were induced similarly by herbivory, independently of previous exposure to HIPVs (Figs. 3g and j).

**Figure 3.**
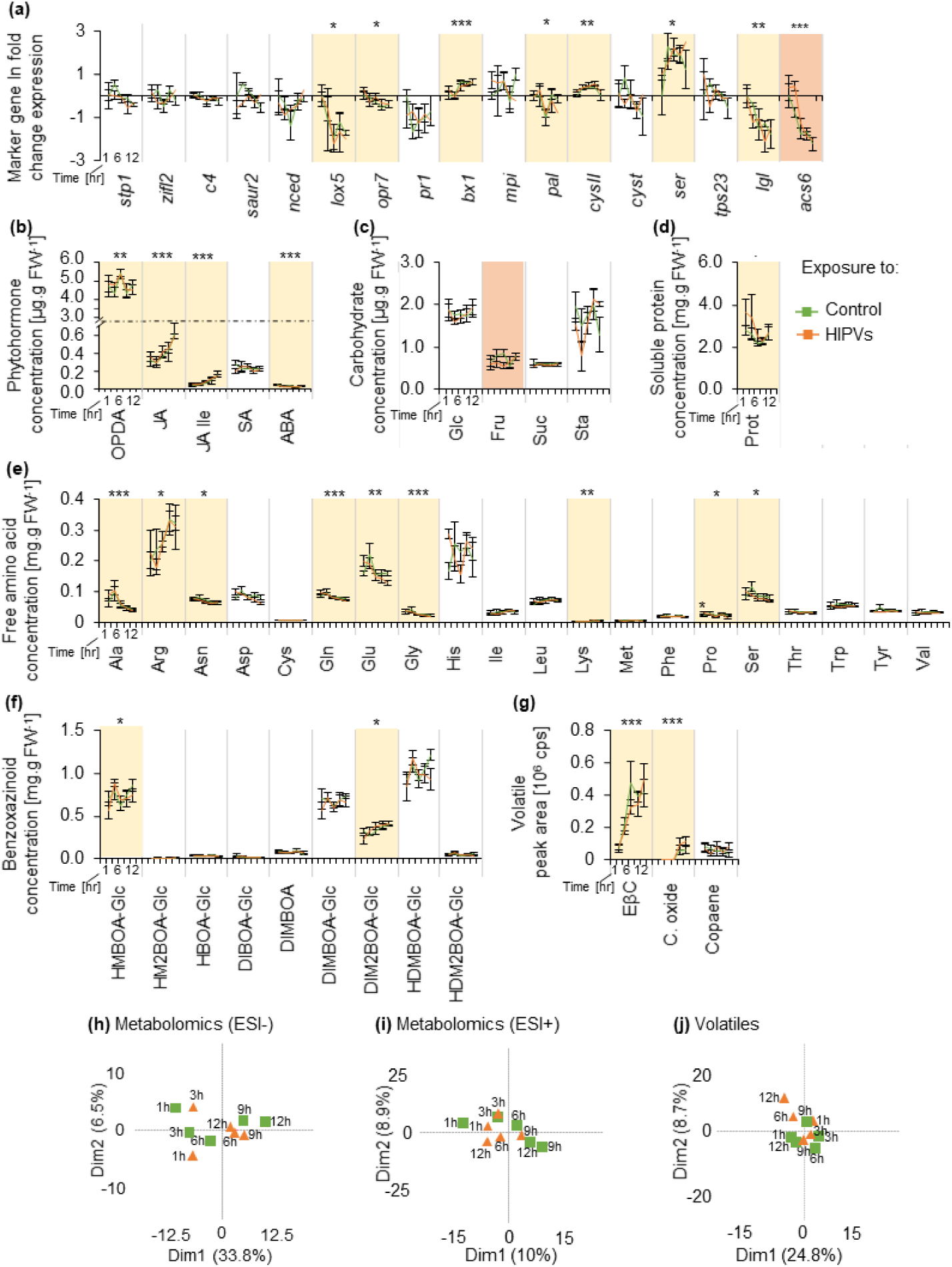
Exposure to an infested neighboring plant does not change the plant response to *D. balteata’s* attack. (**a**) Ln fold changes in gene expression (Mean ± se, Two way ANOVA, n=3-7) in maize roots exposed for four days to plants infested with six *Diabrotica balteata* larvae relative to maize roots exposed to control plants prior attack by *D. balteata* for 1-12 hours. (**b**) Phytohormone production (Mean ± se, Two way ANOVA, n=3-7) maize roots exposed for four days to control plants (control, green) or to plants infested with six *D. balteata* larvae (HIPVs, orange) prior attack by *D. balteata* for 1-12 hours. (**c-f**) Concentrations (Mean ± se, Two way ANOVA, n = 3-7) of (**c**) glucose, fructose, sucrose, and starch, (**d**) proteins, (**e**) amino acids, and (**f**) benzoxazinoids in maize roots exposed for four days to control plants (control, green) or to plants infested with six *D. balteata* larvae (HIPVs, orange) prior attack by *D. balteata* for 1-12 hours. (**h-i**) Principal Component Analysis of all features detected (PPLS DA, n = 3-7) in maize roots exposed for four days to control plants (control, green) or to plants infested with six *D. balteata* larvae (HIPVs, orange) prior attack by *D. balteata* for 1-12 hours, using untargeted metabolomic analysis in (**h**) negative (443 features) and (**i**) positive modes (1906 features). (**j**) Principal Component Analysis of volatile emissions (PPLS DA, n = 3-7) and (**g**) terpene volatiles emissions by maize roots exposed for four days to control plants (control, green) or to plants infested with six *D. balteata* larvae (HIPVs, orange) prior attack by *D. balteata* for 1-12 hours. Only averages per treatment are presented in principal component analyses. EβC: *(E)-*β-caryophyllene. C. oxide: Caryophyllene oxide. Yellow shading and stars indicate significant differences over time (*: p≤ 0·05, **: p ≤ 0·01; ***: p ≤ 0·001). Orange shading indicate significant differences between exposure treatments (p≤ 0·05). No interaction between time and exposure was found to be significant.

### Belowground HIPVs do not increase plant resistance to root herbivory in maize and teosinte

To investigate whether exposure to root HIPVs increases plant resistance in maize or its wild ancestor teosinte, we measured herbivore performance and root damage on control and HIPV-exposed root systems. Exposure to HIPVs emitted by one or three neighboring plants did not alter the herbivore performance, survival, root damage and root fresh mass in both maize and teosinte (Figs. 4, S2).

**Figure 4.**
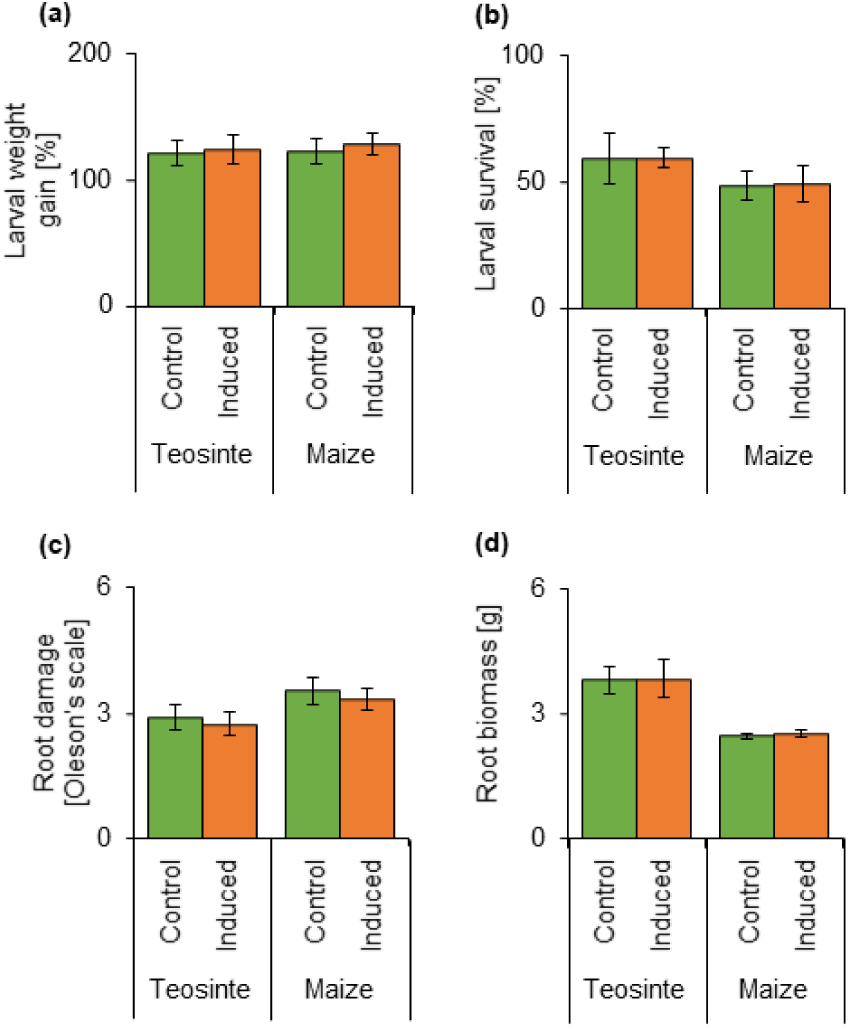
Exposure to an infested neighboring plant does not alter plant defense to herbivory. (**a**) Relative larval weight gain (Mean ± se, Student’s t-tests) of the root herbivore *Diabrotica balteata* feeding for four days on maize (n=17-18) or teosinte (n=8-9) previously exposed for four days to control plants (control, green) or to plants infested with six *D. balteata* larvae (HIPVs, orange). (**b**) Proportions (Mean ± se, Student’s t-tests) of *D. balteata* recovered after 4 days infested on maize (n=18) and teosinte (n=9) previously exposed for four days to control plants (control, green) or to plants infested with six *D. balteata* larvae (HIPVs, orange). (**c**) *D. balteata* damage scaling (Mean ± se, Student’s t-tests) after four days infestation of maize (n=18) and teosinte (n =9) plants previously exposed for four days to control plants (control, green) or to plants infested with six *D. balteata* larvae (HIPVs, orange). (**d**) Root fresh mass after four days infestation by the root herbivore *D. balteata* (Mean ± se, Student’s t-tests) of maize (n=18) and teosinte (n=9) previously exposed for four days to control plants (control, green) or to plants infested with six *D. balteata* larvae (HIPVs, orange).

### Roots are impaired in the emission and perception of resistance-inducing HIPVs

To assess whether roots can perceive and respond to defense-inducing HIPVs, we conducted a cross-experiment where leaf or root tissues were exposed to HIPVs of either leaves or roots prior infestation. Leaf exposure to leaf HIPVs, but not to root HIPVs, lead to a decreased performance of *S. littoralis* caterpillars (Fig. 5a). Root exposure to either leaf or root HIPVs did not affect the root herbivore performance (Fig. 5b). Thus, root HIPVs do not trigger resistance in roots or leaves, and roots, in contrast to leaves, do not respond to leaf HIPVs through an increase in resistance. This result suggests that roots are impaired in both emission and perception of resistance-inducing HIPVs.

**Figure 5.**
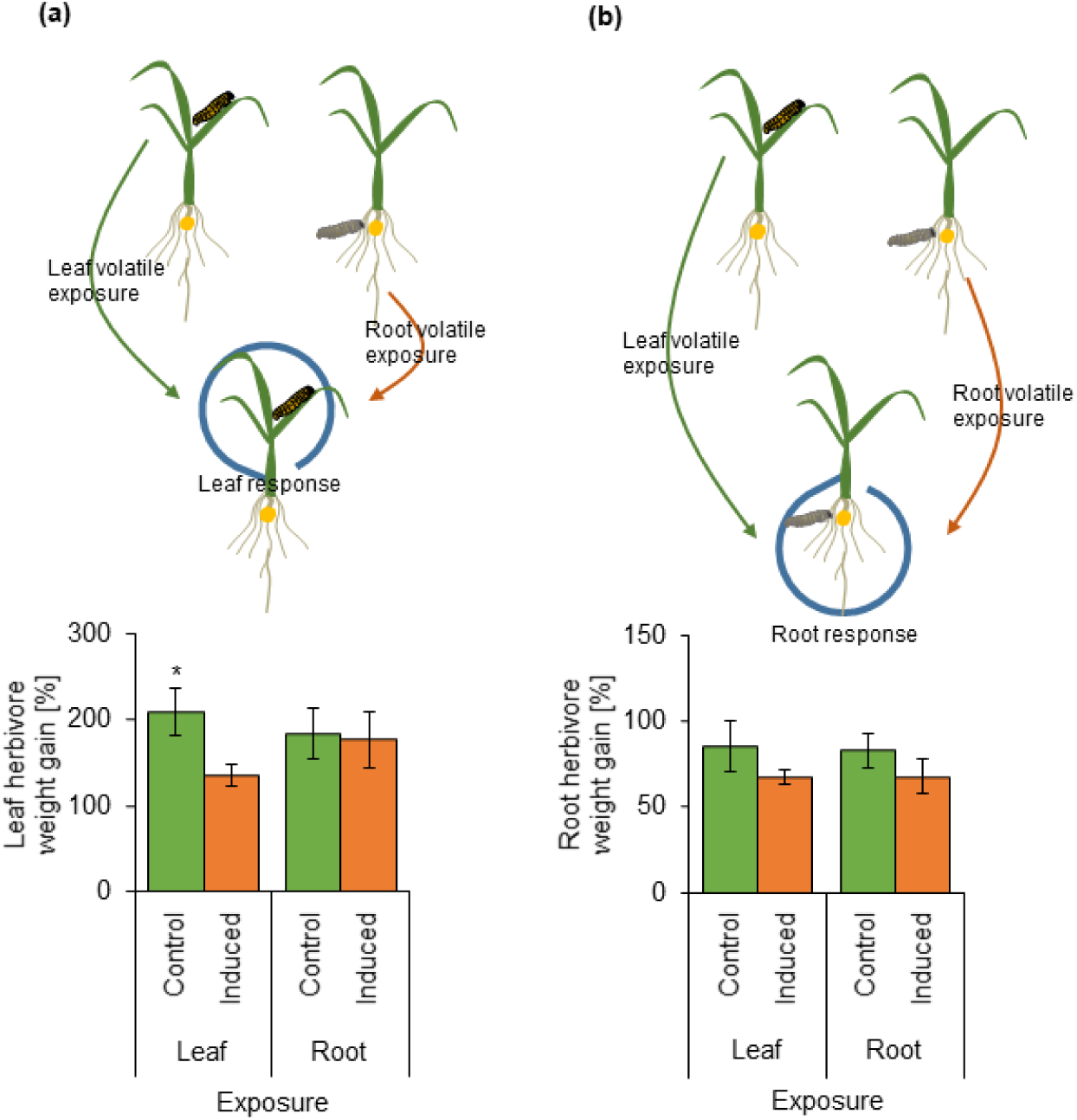
Only leaf exposure to leaf HIPVs leads to a decreased performance of *Spodoptera littoralis* caterpillars. (**a**) Relative larval weight gain (Mean ± se, Two way ANOVA, n=4-5) of the leaf herbivore *S. littoralis* feeding for two days on leaves previously exposed for one night to control plants (control, green) or to plants infested with six *D. balteata* larvae (HIPVs, orange). (**b**) Relative larval weight gain (Mean ± se, Two way ANOVA, n=4-5) of the root herbivore *D. balteata* feeding for two days on roots previously exposed for one night to control plants (control, green) or to plants infested with six *D. balteata* larvae (HIPVs, orange). Stars indicate significant differences within leaf herbivore performance (*: p≤0.05).

## DISCUSSION

The current work shows that HIPV-mediated defense priming occurs in maize leaves, but not roots. The lack of root HIPV response contrasts with the well characterized responses in maize leaves (Engelberth *et al*., 2004; Baldwin *et al*., 2006; Heil & Silva Bueno, 2007; Rodriguez-Saona *et al*., 2009; 2013; Skoczek *et al*., 2017) and is discussed in detail below.

Leaves of many different species are known to respond to HIPVs by increasing their defense investment, and, sometimes also reduce their growth. A recent study furthermore found that volatiles that are constitutively emitted by *Centaurea stoebe* lead to changes in root carbohydrate and protein levels in *Taraxacum officinale* (Gfeller *et al*., 2019; Huang *et al*., 2019). However, *C. stoebe* is an unusually strong constitutive emitter of root terpenes, and whether plants respond to herbivory-induced changes in volatile as a form of “eavesdropping” remains unknown. Our study demonstrates that HIPV-exposed maize roots do not display any of the defense responses displayed by maize leaves and leaves of other plant species (Farmer, 2001; Baldwin *et al*., 2006; Frost *et al*., 2008; Heil & Ton, 2008; Heil, 2014; Erb, 2018; Turlings & Erb, 2018; Bouwmeester *et al*., 2019). Despite prolonged exposure of maize roots to distinct blends of root HIPVs, we did not observe direct induction or priming of stress hormones, primary and secondary metabolites in these roots. On the contrary, we observed that root HIPVs slightly suppressed constitutive JA-Ile levels. This suppression however was gone 1 hr after herbivore attack. Defense marker genes were also not differentially expressed, with the exception of the ethylene biosynthesis gene *acs6*, which was less suppressed upon herbivore attack in HIPV exposed roots. However, these differences were not associated with measurable changes in metabolite accumulation, resistance or plant growth, despite the well-established roles of jasmonates and ethylene in root growth (Staswick *et al*., 1992; Schaller, 2012; Huang *et al*., 2017; Dubois *et al*., 2018) and defense (McConn *et al*., 1997; Bonaventure *et al*., 2011; Erb *et al*., 2012). This absence of phenotypic consequences could be because the changes in Ja-Ile and ethylene biosynthesis were too small and/or transient. Root resistance and plant growth were not affected in teosinte either, suggesting that the absence of HIPV responsiveness in maize roots is not due to plant domestication. From these results, we conclude that maize roots, in contrast to leaves, do not strongly respond to root HIPVs.

What are the physiological mechanisms that could be responsible for the tissue-specific absence of responsiveness of maize roots to root HIPVs? Our experiments suggest two mutually non-exclusive mechanisms: Absence of defense-inducing HIPVs and lack of HIPV responsiveness. Regarding the first mechanism, our experiments show that maize roots do not release any HIPVs that have been shown to mediate priming in maize leaves: GLVs and indole (Farmer, 2001; Engelberth *et al*., 2004; Erb *et al*., 2015; Riedlmeier *et al*., 2017; Ameye *et al*., 2018). Instead, their HIPV profile is dominated by sesquiterpenes (Robert *et al*., 2012a). Sesquiterpenes have been associated with priming in tomato, beans (Arimura *et al*., 2000; Arimura *et al*., 2001; Zhang *et al*., 2019), but not in maize (Ruther & Fürstenau, 2005). This suggests that maize roots do not produce HIPV blends capable of triggering defense responses in neighbors. Why maize roots do not release GLVs and indole remains to be elucidated. GLVs are produced via the hydroperoxide lyase (HPL) branch of the oxylipin pathway (Kenji, 2006). The first step of GLV biosynthesis is to deacylate galactolipids to release the omega-3 and omega-6 fatty acids, α-linolenic acid and linoleic acid (Matsui *et al*., 2000; Kombrink, 2012). The hydroperoxidation of α-linolenic and of linoleic acid results in the production of *Z*-3-hexenal and *n*-hexanal respectively (Moataz *et al*., 2017). Yet, maize roots contains only trace amounts of linolenic acid in favour of high concentrations of linoleic acid (Bernklau & Bjostad, 2008). This limitation in linolenic acid contents in the roots may explain the absence of *Z*-3-hexenal, as well as its alcohol and acetyl GLV downstream products (*Z*-3 and *E*-2 hexenol, *Z*-3 and *E*-2 hexenyl acetate). The lack of indole release is likely due to a different mechanism, as indole-3-glycerol-phosphate, the precursor of indole (Frey *et al*., 2009), is abundant in maize roots. However, the indole-3-glycerol phosphate lyase, which is responsible for volatile indole production (Frey *et al*., 2000) seems to be suppressed upon *D. balteata* attack in the roots, which may explain the absence of volatile indole in the headspace of attacked roots. Regarding the second mechanism, our experiments show that maize roots do not seem capable of increasing their resistance in response to bioactive HIPV blends which are capable of inducing resistance in the leaves. This suggests that maize roots can either not perceive or not translate HIPVs into resistance responses. A better understanding of HIPV perception and early signalling will help to test these hypotheses in the future.

From an adaptive point of view, the question arises why maize plants did evolve the capacity to perceive HIPVs in their leaves, but not their roots. A possible explanation may be that the transfer of HIPVs between plants in the rhizosphere is unreliable. First, volatile dispersal, conversion or degradation in the soil strongly depends on matrix properties (Hayward *et al*., 2001; Owen *et al*., 2007; Perry *et al*., 2007; Hiltpold & Turlings, 2008; Seo *et al*., 2009; Ramirez *et al*., 2010; Peñuelas *et al*., 2014; Xavier *et al*., 2017). Volatile compounds, such as indole, linalool, α-pinene, and limonene, can be degraded upon release and used as source of carbon for soil dwelling micro-organisms (Misra *et al*., 1996; Arora *et al*., 2015; Arora *et al*., 2015; Ma *et al*., 2018; Owen *et al*., 2007; Arora *et al*., 2015; Ma *et al*., 2018). Second, root HIPVs may be less reliable signals, as soil microorganisms produce a wide variety of volatile compounds. Terpenes such as copaene, (*E*)-β-caryophyllene and caryophyllene oxide are also produced by soil micro-organisms (Insam & Seewald, 2010; Wenke *et al*., 2010; Schenkel *et al*., 2015; Delory *et al*., 2016). Thus, we propose that the unreliable transfer and the low specificity of root HIPVs may have impeded the evolution of HIPV perception in maize roots. Instead, alternative strategies to eavesdrop on neighbors may have emerged, including mycorrhizal networks (Perry, 1995; Selosse *et al*., 2006; van der Heijden & Horton, 2009; Jung *et al*., 2012; Song *et al*., 2013; Shahzad *et al*., 2015; Song *et al*., 2019).

In summary, our work shows that plant-plant interactions mediated by herbivore-induced plant volatiles are tissue specific and restricted to the leaves in wild and cultivated maize, and that this tissue-specificity is likely driven by a lack of bioactive cues and a lack of perception capacity of roots. We suggest that the low reliability and specificity of volatiles as danger cues in the rhizosphere together with the availability of other information transfer networks may have impeded the evolution of eavesdropping mechanisms in plant roots.

## ACKNOWLEDGEMENTS

We are really grateful to Anita Streit who reared the insects used in this project. We thank Jean Daniel Berset for his technical assistance. This work was supported by the University of Bern (UniBe 2021) and the Oeschger Centre for Climate Change Research (OCCR).

## AUTHOR CONTRIBUTIONS

CAMR designed the project. CAMR a supervized the project. CvD, TZ, CM, XZ, RARM, RM, MY, BCJS, and GG performed the experiments. CvD, CAMR, TZ, RARM and GG analyzed the data. CvD and CAMR wrote the first draft. All authors reviewed and approved the manuscript.

## Supplementary Information

**Figure S1. The root herbivore *Diabrotica balteata* migrate away from infested plants.** Proportion of larvae escaping from the maize plant after infestation (Mean ± se, One sample t-test, n=6). Stars indicate significant differences (*: p≤ 0·05, **: p ≤ 0·01; ***: p ≤ 0·001).

**Figure S2. Exposure to HIPVs from through infested neighbors does not alter plant defense to herbivory.** (**a**) Relative larval weight gain (Mean ± se, Student’s t-tests) of the root herbivore *Diabrotica balteata* feeding for four days on maize (n=9) previously exposed for four days to control plants (control, green) or to plants infested with six *D. balteata* larvae (HIPVs, orange). (**b**) Proportions (Mean ± se, Student’s t-tests, n=9) of *D. balteata* recovered after 4 days infested on maize previously exposed for four days to control plants (control, green) or to plants infested with six *D. balteata* larvae (HIPVs, orange). (**c**) Root fresh mass after four days exposure to control (green) or to plants infested with six *D. balteata* larvae (HIPVs, orange) and then infested for four days by the root herbivore *D. balteata* (Mean ± se, Student’s t-tests, n=9).

**Table S1. Primer list for q-RT-PCR used to assess the plant response in this study** (Peng *et al*., 2005; Ton *et al*., 2007; Gao *et al*., 2008; Erb *et al*., 2009; Robert *et al*., 2012b; Remy *et al*., 2014; Hajiahmadi *et al*., 2017); *NCBI Gene: 100193700*^*^).

