## Supplementary Table for "Tissue-specific volatile-mediated defense regulation in maize leaves and roots"

**Table S1. Primer list for q-RT-PCR used to assess the plant response in this study** (Peng *et al.*, 2005; Ton *et al.*, 2007; Gao *et al.*, 2008; Erb *et al.*, 2009; Robert *et al.*, 2012b; Remy *et al.*, 2014; Hajiahmadi *et al.*, 2017); *NCBI Gene:* [*100193700*](https://www.ncbi.nlm.nih.gov/sites/entrez?db=gene&cmd=search&term=100193700)^*^)

| Gene name | Putative function | Forward primer (3’-5’) | Reverse primer (5’-3’) |
| --- | --- | --- | --- |
| *Zm-actin1* | Actin | ccatgaggccacgtacaact | ggtaaaaccccactgagga |
| *Zm-gapc* | Glyceraldehyde phosphate dehydrogenase | gcatcaggaaccctgaggaa | catgggtgcatctttgcttg |
| *Zm-cysII* | Cystatin II proteinase inhibitor | tgccctgctcatactgcttg | gcgagttcctggaggtgaag |
| *Zm-cyst.* | Cystatin proteinase inhibitor | caaggagcacaacaggcaga | ggacatgagctggcgatttt |
| *Zm-bx1* | Indole-3-glycerol phosphate lyase | cccgagcacgtaaagcagat | cttcatgcccctggcatact |
| *Zm-igl* | Indole-3-glycerol phosphate lyase | gcctcatagttcccgacctc | gaatcctcgtgaagctcgtg |
| *Zm-nced* | ABA biosynthesis | aagtctaccatccacacagg | tcaacggagccagctgat |
| *Zm-opr7* | Jasmonic acid synthase | cggctgttcatcgctaatcccgc | caatcgcggcattacccagatgt |
| *Zm-pr1* | Pathogenesis-related gene 1 | ctgggtgtccgagaagcagt | cgggttgtagctgcagatgat |
| *Zm-saur2* | Auxin biosynthesis gene | gtgccttagcacccctgtct | ggctcctctcctgagcaaac |
| *Zm-serpin* | Serine proteinase inhibitor | gacggaggaggaaggaggag | acctgatgcactgcttgcac |
| *Zm-stp1^*^* | Carbohydrate transporter | ttcgccaaccagtccgtgcc | cagccgcccctgatcttggc |
| *Zm-zifl2* | Carbohydrate transmembrane transporter | gggagccactgctggcgaag | cggcagggtgcaggtgatgg |
| *Zm-mpi* | Proteinase inhibitor | aagctttttaggttctacacaaaaccctc | tctagaccggaccagttgacga |
| *Zm-pal* | Phenylalanine  ammonia lyase | cgaggtcaactccgtgaacg | gctctgcacgtggttggtga |
| *Zm-lox5* | Lipoxygenases | gcggtgatcgagccgttcgtaatc | acggcgcagcacaacaatttactacgag |
| *Zm-acs6* | Ethylene synthase | ctggatcaacctcggctaacga | tggctagcaaatgttgagtttg |
| *Zm-tps23* | Terpene synthase | tctggatgatgggagtcttctttg | gcgttgccttcctctgtgg |
