## Supplementary Figures for "Tissue-specific volatile-mediated defense regulation in maize leaves and roots"

### Slide 1
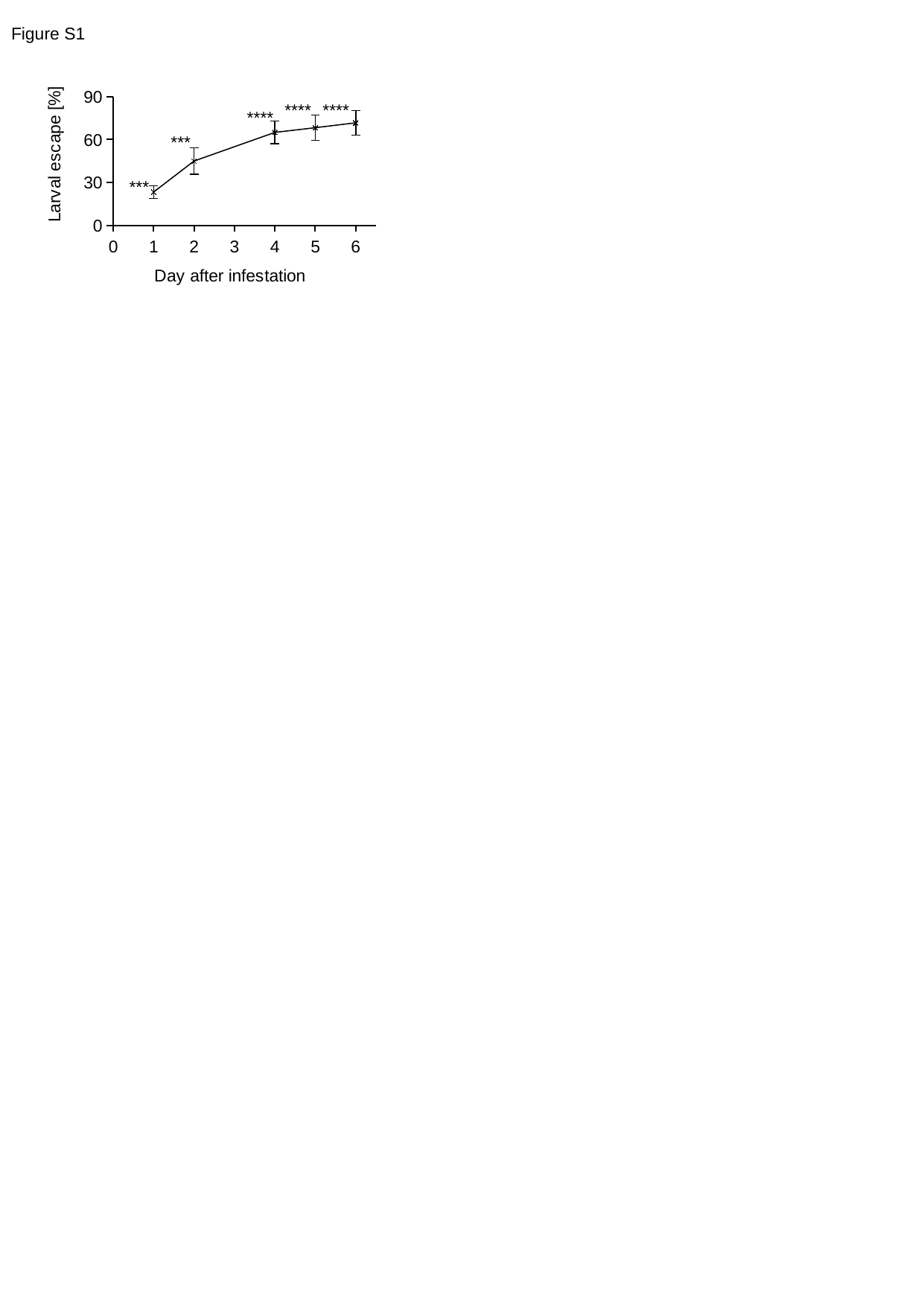

Figure S1
#### Chart
| Category | |
|---|---|****
****
****
***
***

### Slide 2
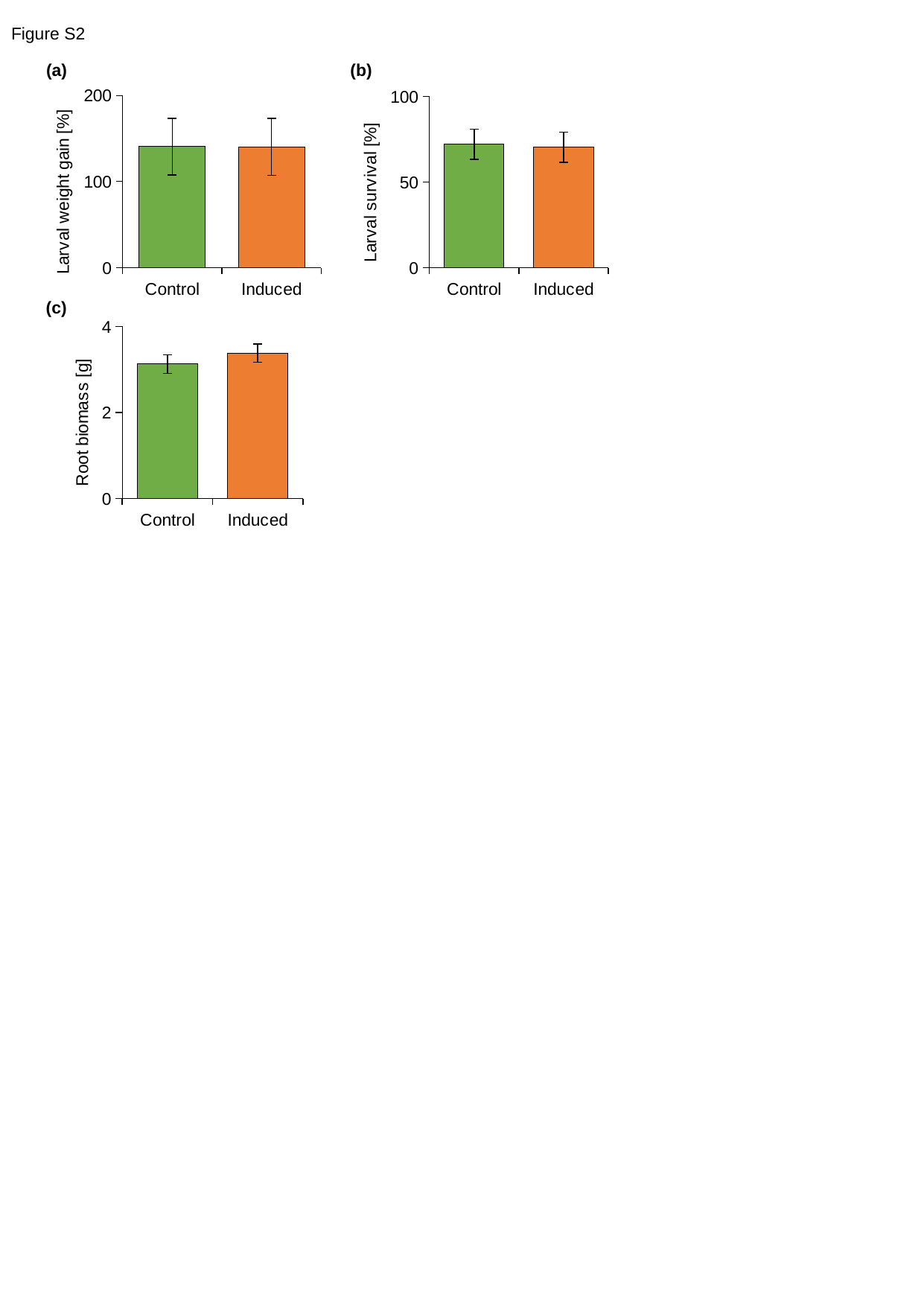

Figure S2
(a)
(b)
#### Chart
| Category | gain weigh (%) |
|---|---|
| Control | 140.71210991727264 |
| Induced | 140.5499979703758 |
#### Chart
| Category | Recver (%) |
|---|---|
| Control | 72.22222222222223 |
| Induced | 70.37037037037037 |
#### Chart
| Category | total root weight (mg) |
|---|---|
| Control | 3.125777777777778 |
| Induced | 3.38 |(c)
